# Double-transmembrane domain of SNAREs decelerates the fusion by increasing the protein-lipid mismatch

**DOI:** 10.1101/2020.08.24.264663

**Authors:** Bing Bu, Zhiqi Tian, Dechang Li, Kai Zhang, Wei Chen, Baohua Ji, Jiajie Diao

## Abstract

SNARE is the essential mediator of membrane fusion that highly relies on the molecular structure of SNAREs. For instance, the protein syntaxin-1 involving in neuronal SNAREs, has a single transmembrane domain (sTMD) leading to fast fusion, while the syntaxin 17 has a V-shape double TMDs (dTMDs), taking part in the autophagosome maturation. However, it is not clear how the TMD structure influences the fusion process. Here, we demonstrate that the dTMDs significantly reduce fusion rate compared with the sTMD by using an in vitro reconstitution system. Through theoretical analysis, we reveal that the V-shape dTMDs can significantly increase protein-lipid mismatch, thereby raising the energy barrier of the fusion, and that increasing the number of SNAREs can reduce the energy barrier or protein-lipid mismatch. This study provides a physical-chemical mechanistic understanding of SNARE-regulated membrane fusion.

## INTRODUCTION

As a key cellular process, membrane fusion plays a decisive role in the neurotransmission, drug delivery, exo- and endocytosis [1-4]. Soluble N-ethylmaleimide-sensitive factor activating protein receptors (SNAREs) serve as the molecular machine to mediate neurotransmission and other fusion process [5, 6]. The core structure of neuronal SNAREs is composed of synaptobrevin-2 (Syb 2, also called VAMP2: vesicle-associated membrane protein 2), syntaxin-1 (Syx 1), and SNAP-25. The C-terminal single transmembrane domain (TMD) is one of the key structures of the SNAREs for the fusion. The TMD domain in Syb 2 anchored on the synaptic vesicles, while that in Syx 1 located in the plasma membrane. They winded with SNAP-25 and formed a 4-helical SNARE core structure [7]. The extension of the zipper formation of the core structure to membranes by these two single TMDs is the main driving force of membrane fusion [8]. Most of the SNARE proteins have single TMD (sTMD), however, syntaxin 17 (Syx 17) serving in the fusion between autophagosomes and lysosomes contains the V-shape double TMDs (dTMDs) [9, 10] (**Fig. S1**). It is interesting to find the roles of Syx 17 dTMDs in membrane fusion for the mechanistic understanding of fusion regulation.

The TMD and its interaction with membrane influences the process of membrane fusion, which is determined by the physicochemical property of the lipids and the structure of the TMD [1, 11-16]. For instance, Katsov et al. [11] investigated how lipid compositions influenced fusion process, suggesting that lipids with spontaneous negative curvature (e.g. phosphatidylethanolamine (PE) and cholesterol (Chol)) have a lower energy barrier to accomplish fusion process. Furthermore, Jackson took into account the deformation and motion of the membrane and SNAREs TMDs into a theoretical model and predicted the effect of TMD properties on the profile of fusion energy and the fusion rate [13], the number and stiffness/flexibility of TMDs can regulate fusion process and kinetics [13, 17]. However, previous studies mainly focused on either the SNAREs or the membrane individually, the effect of SNAREs-lipid interaction on the fusion process is still unclear.

When the TMD domain of SNAREs (i.e. Syb and Syx) insert into the membrane, a protein-lipid mismatch may exist between the TMDs and the lipid bilayer due to the difference between the length of TMDs and the hydrophobic thickness of the lipid bilayer [18-22]. Such a protein-lipid mismatch can tilt the insertion angle of the TMDs depending on the length and sequence of TMDs and the membrane composition [18-20]. Therefore, this protein-lipid mismatch can change the interaction and local distribution of SNAREs and thus influence the fusion rate [23, 24]. To date, however, there is very few models considering the role of the SNAREs-lipid interaction in regulating the membrane fusion process. Motivated by these results, we proposed that SNAREs-lipid interaction should play critical roles in the membrane fusion.

To study the role of TMD structure in membrane fusion, we developed an approach of changing TMD, different from previous assay of removing TMD [25]. Through ensemble lipid-mixing and single-vesicle docking assays, we found that the dTMDs of Syx 17 reduced the fusion rate about 4 times compared to the sTMD domain from the wildtype Syx 1 (Syx 1 WT). To explain this difference, we analyzed the effect of SNAREs-lipid mismatch on the membrane fusion process by introducing a theoretical model that treats the protein-lipid interaction explicitly. This model predicted that the increased protein-lipid mismatch by the dTMDs slowed down the fusion process, which provides the theoretical explanation of the underlying mechanisms of the experimental results. In addition, this study built both theoretical and experimental frameworks for the study of the regulatory roles of protein-lipid interaction involved in the membrane fusion process.

## MATERIALS AND METHODS

### Protein preparation

All SNARE proteins were from rat and expressed and purified as described by [26-28]. Briefly, his-tagged Syx 1 WT, Syx 1/17 containing the cytoplasmic domain of Syx 1 and the dTMDs of Sxy 17, Syb 2, and SNAP-25 were expressed in *E. coli* and purified using a combination of Ni-NTA affinity (Qiagen, Hilden, Germany) and size exclusion chromatography on a Superdex 200 column (GE Healthcare, Uppsala, Sweden). His-tags were removed with TEV protease.

### Ensemble lipid mixing

A step-by-step protocol for v-/t-SNARE vesicle reconstitution for lipid mixing experiments has been reported in our previous publication [29]. The protein to lipid ratio was 1:200 for both t-SNARE and v-SNARE vesicles, by which approximately 100-200 copies of SNARE proteins would be reconstituted to individual vesicles [30, 31]. The lipid composition was 2:12:20:20:46 = DiI(DiD):PS:PE:Chol:PC. DiI-labeled t-SNARE vesicles and DiD-labeled v-SNARE vesicles were mixed at a molar ratio of 1:1. To demonstrate the activity of vesicle fusion via lipid mixing, we measured acceptor fluorescence intensity by FRET using a fluorescence spectrometer (Varian Cary). Wavelengths of 530 and 670 nm were used for excitation of donor (DiI) and emission of acceptor (DiD), respectively. All experiments were performed at 35 °C.

### Single vesicle docking

A detailed protocol for this step has been previously described [32, 33]. The PEGylated surface of the microfluidic chamber was incubated with neutravidin (Invitrogen) for 5 min and washed with buffer.

The v-vesicles were immobilized on the surface with a 5-min incubation, and washed with buffer to remove free vesicles. Then, t-vesicles were injected and washed after 15 min of incubation. Docked t-vesicles were excited by a 532-nm laser (Crystal laser) on a total internal reflection fluorescence microscopy (Nikon). The docking number per an imaging area (45 × 90 μm^2^) was analyzed and averaged by using a customized program written in C++ (Microsoft).

## RESULTS

### Effect of dTMDs on the fusion rate

To check the influence of dTMDs on fusion, we first performed experiments to investigate how the protein-lipid mismatch changes the fusion rate. Firstly, the dTMDs of Syx 17 were hybridized with Syx 1 WT (named as Syx 1/17) to eliminate the residue sequences difference of their zipping domains. We then performed an ensemble lipid-mixing assay to study the influence of dTMDs on the fusion process (**Fig. 1A**). The fluorescence intensity produced by FRET between the donor and acceptor dyes in vesicles was measured for ∼1800 s (**Fig. 1B**). The same v-SNARE vesicles reconstituted with Syb 2 were used for vesicles reconstituted with Syx 1 WT (with sTMDs) and Syx 1/17 (with dTMDs). During the whole timecourse of fusion, the intensity of Syx 1 WT system is higher than that of Syx 1/17, indicating that the fusion rate of Syx 1 WT system is higher than that of Syx 1/17. By fitting the fluorescence intensity curve, we found that the fusion rate K of Syx 1 WT system is ∼3.6-4.1 times as much as that of Syx 1/17. To eliminate the influence of dTMDs on vesicle docking, we also performed the single-vesicle docking assay (**Fig. S2**). No difference on docking was observed (**Fig. S2B**), indicating that replacing TMD of Syx 1 has little influence on the early stages of SNAREs zipping [34]; and the reduced lipid mixing lies on the fusion step. Moreover, since the vesicle docking is induced by the interaction of SNARE N-terminal domains, the result shown in **Fig. S2B** also implies the reconstituted level of t-SNARE proteins for Syx 1 WT and Syx 1/17 vesicles was similar before TMDs comes to close contact [8] and the fusion reduction was mainly caused by the structural difference between dTMDs and sTMDs.

**Figure 1.**
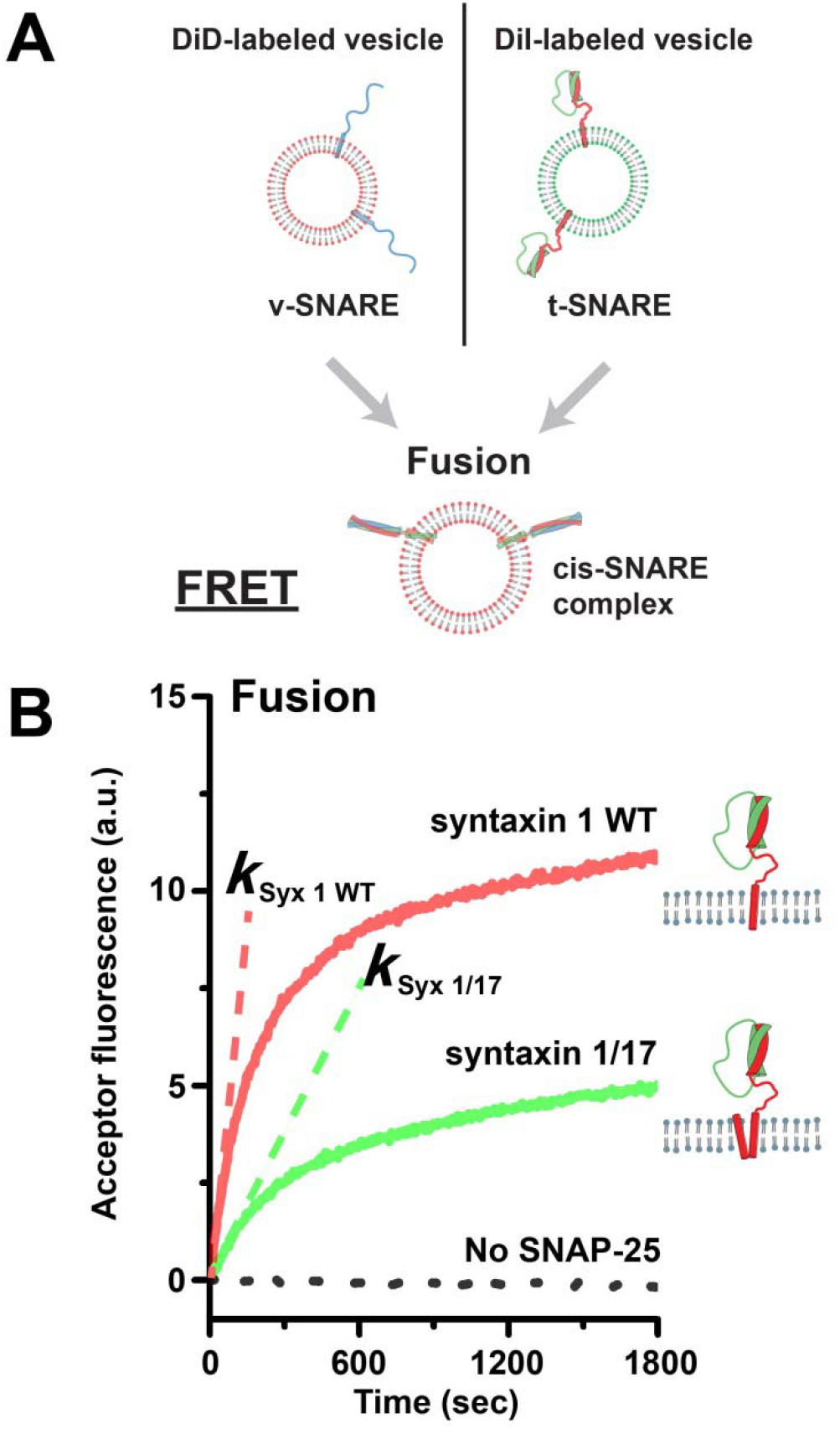
dTMDs reduce the rate of membrane fusion in vitro. (A) The illustration of v-vesicles and t-vesicles fusion. (B) Fusion of v-vesicles and t-vesicles reconstituted with syntaxin 1 wild-type (Syx 1 WT) with sTMDs and syntaxin 1 hybridized to the dTMDs of syntaxin 17 (Syx 1/17). The y axis is the acceptor fluorescence intensity produced by FRET between the donor and acceptor dyes in vesicles, a measure of the activity of vesicle fusion with lipid mixing.

### Theoretical estimation of fusion energy

A theoretical model was introduced to investigate the effect of protein-lipid mismatch between the length of TMDs and the membrane thickness on the membrane fusion. Three representative structures were used to capture the fusion process (**Fig. 2A)** [35]. In the beginning, the tilted TMDs of Syx and Syb formed bundles in the corresponding membranes (**Fig. 2A**, *α* state); with the zipping of the SNAREs core helical structure, the TMDs rotated and moved along with the deformation of the membrane (**Fig. 2A**, *β* state); when the SNARE zipping finished, the two opposed membranes merged and a fusion pore formed, the TMDs of Syx and Syb came to close contact (**Fig. 2A**, *γ* state). In the theoretical model, we described the fusion process with a reaction coordinate *d*, the distance between the tails of transmembrane domain, as shown in the state *β* of **Fig. 2A**. This distance is at its minimum, *d*_min_, at the state *α* ; and it reaches its maximum, *d*_max_ = 2*b*, after fusion at the state of *γ*, where *b* is the length of TMDs. Because *d* monotonically increases from *α* to *γ* state, we use its normalized form, *ξ* (*d*) = (*d* − *d*_min_) (*d*_max_ − *d*_min_) as the fusion coordinate, to describe the fusion process. Note that *ξ* (*d*) = 0 at state *α* and *ξ* (*d*) = 1 at state *γ*. The energy involved in the fusion process was divided into three parts

**Figure 2.**
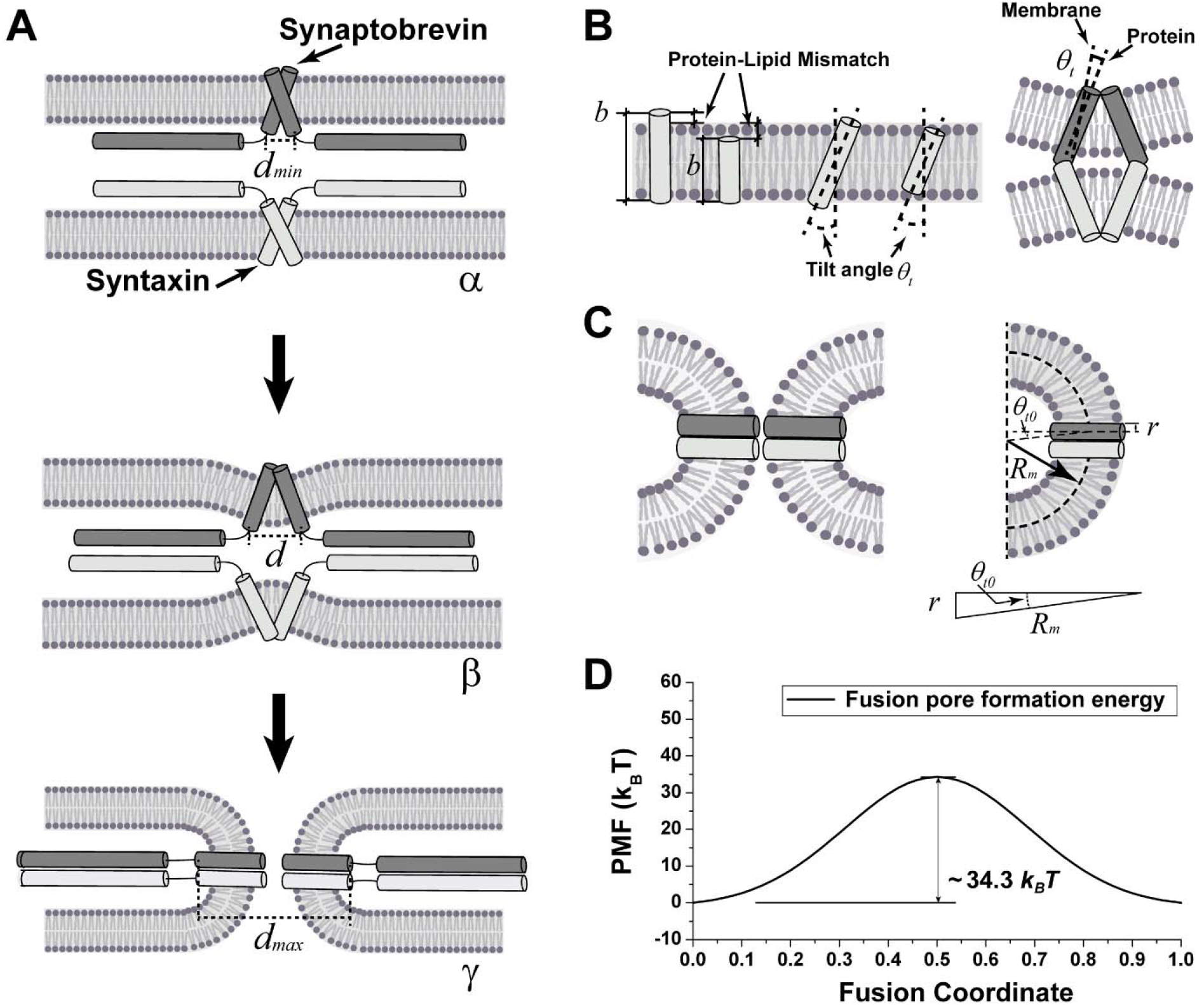
The fusion process with protein-lipid mismatch. (A) The representative structures of transition process from protein anchored in the membrane to fusion pore formation. TMDs rotated and moved, membrane deformed with fusion process from *α* to *γ*. *d*_min_, *d*, and *d*_max_ is the distance between the TMD’s tails at different states, respectively. (B) The illustration of *E*_*t*_ energy contribution. *E*_*t*_ is the energy caused by the TMDs tilt (protein-lipid mismatch). *b* is the length of TMDs. *θ*_*t*_ is the tilt angle between the direction of TMDs and normal direction of membrane. (C) Illustration of protein-lipid mismatch interface in *γ* state. *θ*_*t*0_ is the residual tilt angle when the fusion process finished. *R*_*m*_ is the radius of curvature of membrane contour. *r* is the radius of TMD. The residual tilt angle *θ*_*t*0_ can be calculated as *θ*_*t*0_ = arcsin (*r R*_*m*_). (D) The profile of fusion pore formation energy *E*_*pore*_ as a function of the fusion process was set as a Gauss curvature [46].

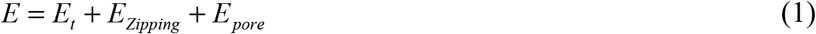

in which *E*_*t*_ is the energy induced by the protein-lipid mismatch; *E*_*Zipping*_ is the energy released by the SNARE zipping; and *E*_*pore*_ is the energy cost of the membrane deformation for forming a fusion pore.

The protein-lipid mismatch may exist due to the difference between the hydrophobic thickness of lipid bilayer and the length of TMDs [18-22]. For TMDs with positive or negative protein-lipid mismatch, the hydrophobic mismatch or rotation entropy will be induced when the TMDs insert into the membrane vertically [19, 20, 36] (**Fig. 2B**). Thus, the TMDs prefer to insert into the membrane with a suitable tilt angle 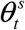 to accommodate the mismatch between TMDs and the lipid bilayer (**Fig. 2B**) [19, 20]. Accordingly, the total interfacial energy depends on the tilt angle. Here we assume that the tile energy *E*_*t*_ changes linearly with number of the TMDs in the membrane during the fusion process (**Fig. 2A** *α* to *γ* state), therefore the tilt energy is given by

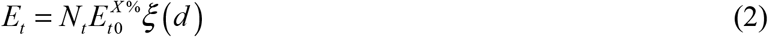

where *N*_*t*_ is the number of TMDs involving in the fusion process. For single SNAREs complex, *N*_*t*_ = 2 (TMDs from Syx1 WT and Syb). For two SNAREs complex, *N*_*t*_ = 4 etc. 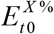 is the change of the tilt energy for one TMD during the fusion process. Previous studies [19, 20] showed that the tilt energy 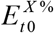 depends on the length of TMDs and the hydrophobic thickness of the membrane [18, 20, 22], *X* % is a label for membrane composition contains *X* % Chol, which will be described later. As shown in **Fig. 2C**, when the fusion process finishes (**Fig. 2A**, *γ* state) and Syx comes to closely contact with Syb, both the TMDs insert in the membrane with a residual tilt angle 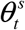 due to the radius of TMDs (**Fig. 2C**). The radius of membrane contour curvature *R*_*m*_ (**Fig. 2C**) was estimated in a range of 3-10 nm based on previous simulations and experiment results [13, 37-41], the radius of TMDs is *r* = 0.35*nm* [13]. Therefore, the residual tilt angle when the fusion process finishes can be calculated by *θ*_*t*0_ = arcsin (*r R*_*m*_) (**Fig. 2C**), and 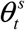 comes to be ∼2.0 − 6.7°. During the fusion process, the tilt angle changes from the suitable tilt angle 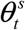 to *θ*_*t*0_. For a membrane with the hydrophobic thickness ∼2.68 nm and the TMD length ∼3.15 nm, the suitable title angle is 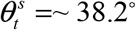 and the tilt energy is 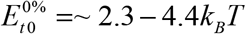 according to the umbrella sampling calculations [20].

Previous studies showed that the hydrophobic thickness of the membrane can be regulated by the proportion of Chol [42, 43]. It was shown that Chol can increase the hydrophobic thickness of the membrane by ∼0.32 nm with ∼29% proportion and the increase is linear to the Chol proportion [42]. For a membrane with pure DOPC, the hydrophobic thickness is ∼2.68 nm [44], while the TMD of Syx contains 21 residues which has a length of ∼3.15 nm. When the Chol proportion reaches to ∼42.6%, the hydrophobic thickness of the membrane will be close to 3.15 nm, and the protein-lipid mismatch will be negligible. Note that although Chol can reduce the hydrophobic length mismatch between lipid and TMD, the tilt angle will still exist even if the hydrophobic length mismatch is negligible [20]. For example, the TMD may have a ∼ 10o inherent tilt angle and considerable helix rotation which arise from the helix precession around the membrane normal [19, 20, 36]. When the hydrophobic thickness of membrane is equal to the length of TMD, the tilt energy 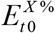 is attributed to the rotational entropy contribution of the helix precession around the membrane normal, calculating as 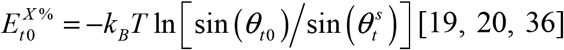. So that the change of the tilt energy with 42.6% Chol during the fusion process 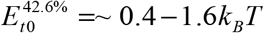 when the hydrophobic thickness of membrane is equal to the length of TMD. We assumed that the tilt energy 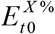 depends on the Chol proportion linearly. So that for 20% Chol proportions, 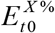 comes to be 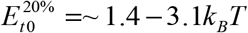, respectively.

*E*_*Zipping*_ is the energy of SNAREs zipping which are assumed to be linear with {(d)

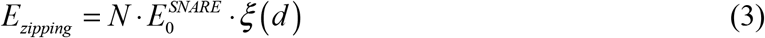

where 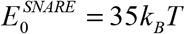 is the energy of each SNARE released during the fusion process [45].

Finally, *E*_*pore*_ is the energy cost for creating a fusion pore on the membrane during the fusion process. The membrane deformation during fusion process should pass several metastable states and overcome energy barriers for, e.g., stalk, hemifusion, and fusion pore formation. These energy barriers can be represented by one effective energy barrier [46] and simplified as a Gaussian-like form [47]. The energy profile changing with the fusion coordinate is fitted with an energy barrier of 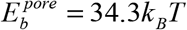 [46] as

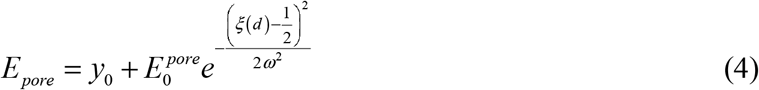

The parameters used in this study were listed in **Table 1**.

**Table 1.** The definitions and parameters for the theoretical model.

| Symbol | Definition | Values | Ref |
| --- | --- | --- | --- |
| $N$ | Number of SNARE complexes | 1-3 | [50] |
| $r$ | Radius of TMDs | $3.5\text{ \AA}$ | [13] |
| $R_m$ | Radius of membrane contour curvature | 3-10nm | [13, 37-41] |
| $E_{t0}^{0\%}$ | Tilt energy change during fusion process with 0% Chol | $\sim 2.3-4.4k_BT$ | [20] |
| $E_b^{pore}$ | Energy to generate a membrane fusion pore | $34.3k_BT$ | [46] |
| $y_0$ | Gauss curvature parameter | $-1.01\text{ }k_BT$ | Fitting with [46] |
| $E_0^{pore}$ | Gauss curvature parameter | $35.3\text{ }k_BT$ | Fitting with [46] |
| $\omega$ | Gauss curvature parameter | 0.188 | Fitting with [46] |
| $E_{SNARE}$ | Energy provided by one SNARE protein | $35k_BT$ | [45] |

To calculate the fusion rate with the Kramers’ theory [13, 48, 49], the reaction rate *k* can be written as [13]

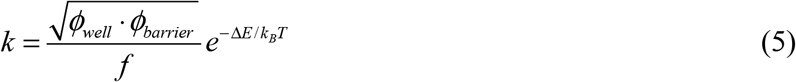

where Δ*E* is the energy barrier of the reaction. *Φ*_*well*_ and *ϕ*_*barrier*_ are the quadratic coefficients for the energy profile at the minimum and at the peak of the barrier, respectively. *f* is a constant related to the diffusion coefficient. *k*_*B*_ is the Boltzmann constant, and *T* is the absolute temperature. Finally, we got the ratio of the fusion rate between two reactions of *i* and *j*

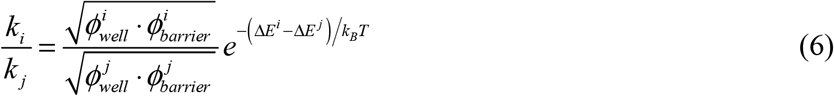

### Effect of protein-lipid mismatch

The total energy of membrane fusion was calculated for different number of SNAREs. The energy profiles and energy barrier for N = 1∼3 were shown in **Fig. 3A**. Our data show that the energy barriers decrease with the increasing of the number of SNAREs taking part in the fusion process. When N = 3, the energy barrier for the fusion nearly vanishes (see **Fig. 3A**). These data suggest that the fusion is highly accelerated when more SNAREs are involved, consistent with previous study showing that efficient fusion requires three or more SNARE complexes [50].

**Figure 3.**
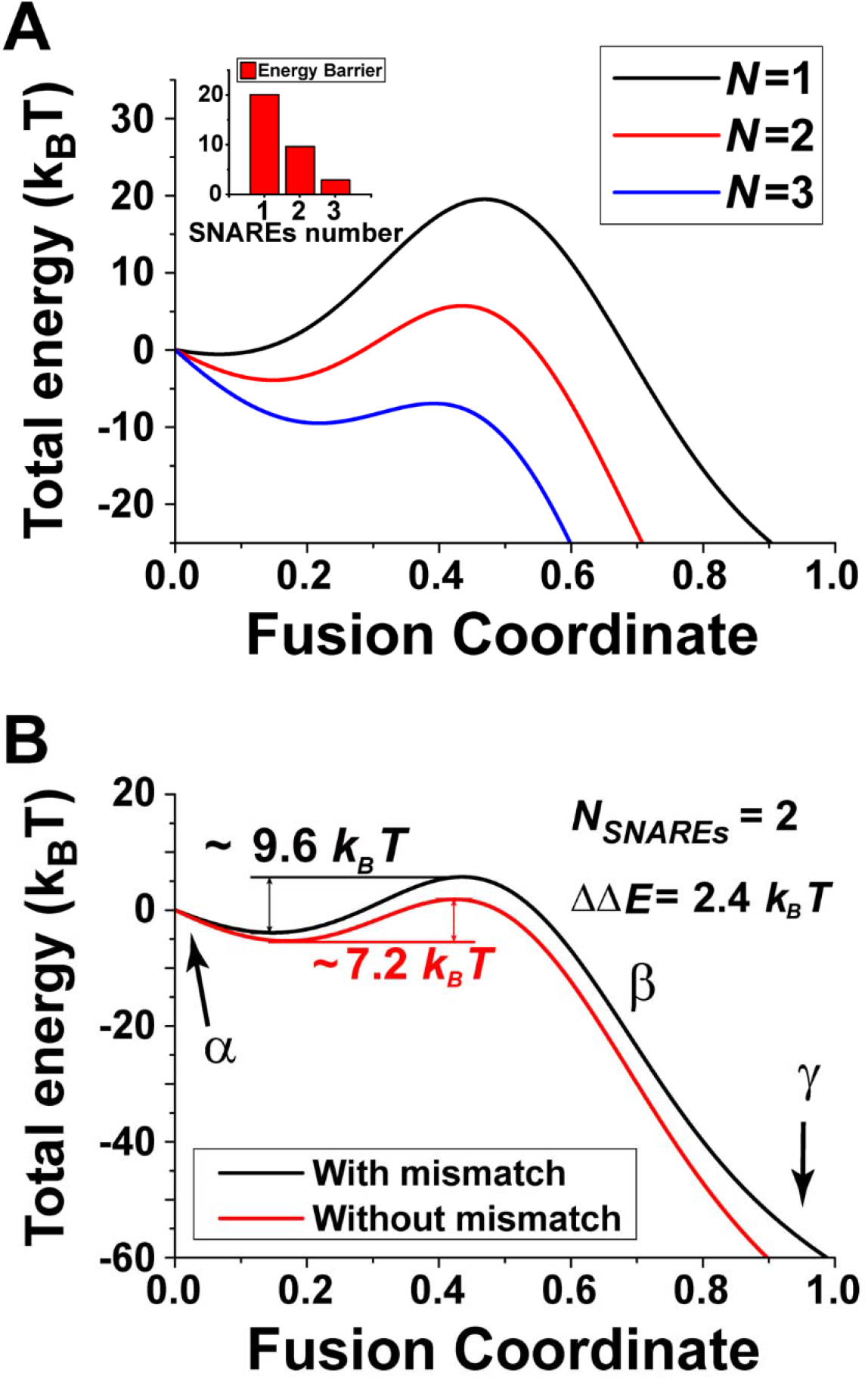
(A) The fusion energy profiles and barrier changed with different SNAREs number. (B) The energy profile of fusion process with and without mismatch as a function of fusion coordinate. Two SNAREs were assumed to take part in the fusion process. To consistent with the experiment, the *E*_t0_ = 2.25*k*_*B*_*T* was used in the theoretical claculation with 20% Chol.

To investigate the influence of protein-lipid mismatch between the TMDs and lipid bilayer on the fusion process, we calculated the energy profile with two SNAREs during fusion in the presence and absence of protein-lipid mismatch. The energy barrier of fusion process in the presence of protein-lipid mismatch is ∼2.4*k*_*B*_*T* higher (**Fig. 3B**). At the beginning of the protein-lipid-mismatch fusion, the TMDs are tilted in the membrane (see *α* state in **Fig. 2A**). After the formation of fusion pore, the TMDs of Syb and Syx contact with each other and are almost perpendicular to the membrane (see *γ* state in **Fig. 2A** and **Fig. 2C**). It is clearly suggested that the increase of energy barrier is caused by the protein-lipid mismatch. Thus, the fusion process with the protein-lipid mismatch experiences a higher energy barrier, leading to a slower fusion rate.

### Effect of SNARE number

To further understand our experimental results showing double-TMDs reduce fusion (**Fig. 1**), we analyzed the influence of dTMDs with our theoretical model. The change of the tilt energy during the fusion process with dTMDs is larger than that with sTMD because of the increase of TMD numbers *N*_*t*_. The difference in the geometric arrangement of TMDs and their interaction with lipid bilayer between the systems with sTMD and dTMDs is illustrated in **Fig. S3**. The energy profile during the fusion process is shown in **Fig. 4A**. And we showed that, when 1-3 SNAREs were involved in the fusion process, the fusion rate of Syx 1 WT could be ∼1.7-7.7 times higher than that of Syx 1/17 (**Fig. 4B**). For example, when two SNAREs take part in the fusion process, the dTMDs increase the fusion energy barrier and reduces the fusion rate. The energy barrier of Syx 1/17 is ∼1.33*k*_*B*_*T* higher than that of Syx 1 WT, so that the fusion rate of Syx1 WT is ∼3.8 times higher than that of Syx 1/17, consistent with the experimental results (**Fig. 1B**). Note that various experiment methods get different SNAREs energy 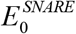 it is ∼ 68 − 85*k*_*B*_*T* with magnetic and optical tweezers [34, 51]. Besides, the estimates obtained with atomic-force microscopy or the surface-force apparatus are much lower, which was in the 30*k*_*B*_*T* range [45, 51]. Li et al. suggested that SNAREs release 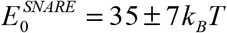 [45]. Considering the error effect of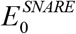, we calculated *k*_*Syx* 1 *WT /*_*k*_*Syx* 1/17_ when 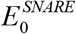 varies from 28 to 42 *k*_*B*_*T* with tilt energy *E*_t0_ = 2.25*k T* (20% Chol). When 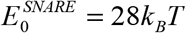, the fusion rate of Syx 1 WT could be ∼2.6-6.2 times higher than that of Syx 1/17; when 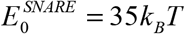, the fusion rate of Syx 1 WT could be ∼2.5-3.9 times higher than that of Syx 1/17; when 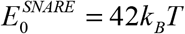, the fusion rate of Syx 1 WT could be ∼2.3-3.9 times higher than that of Syx 1/17. Both our experimental and theoretical results show that the dTMDs reduce fusion rate significantly, which can be explained why the fusion process of autophagosomes and lysosomes is slower than the fusion in neurotransmission [25, 52]. In addition, our results suggest that the number of SNAREs involved in a general fusion process is likely to be 2 to 3 (see **Fig. 4B**).

**Figure 4.**
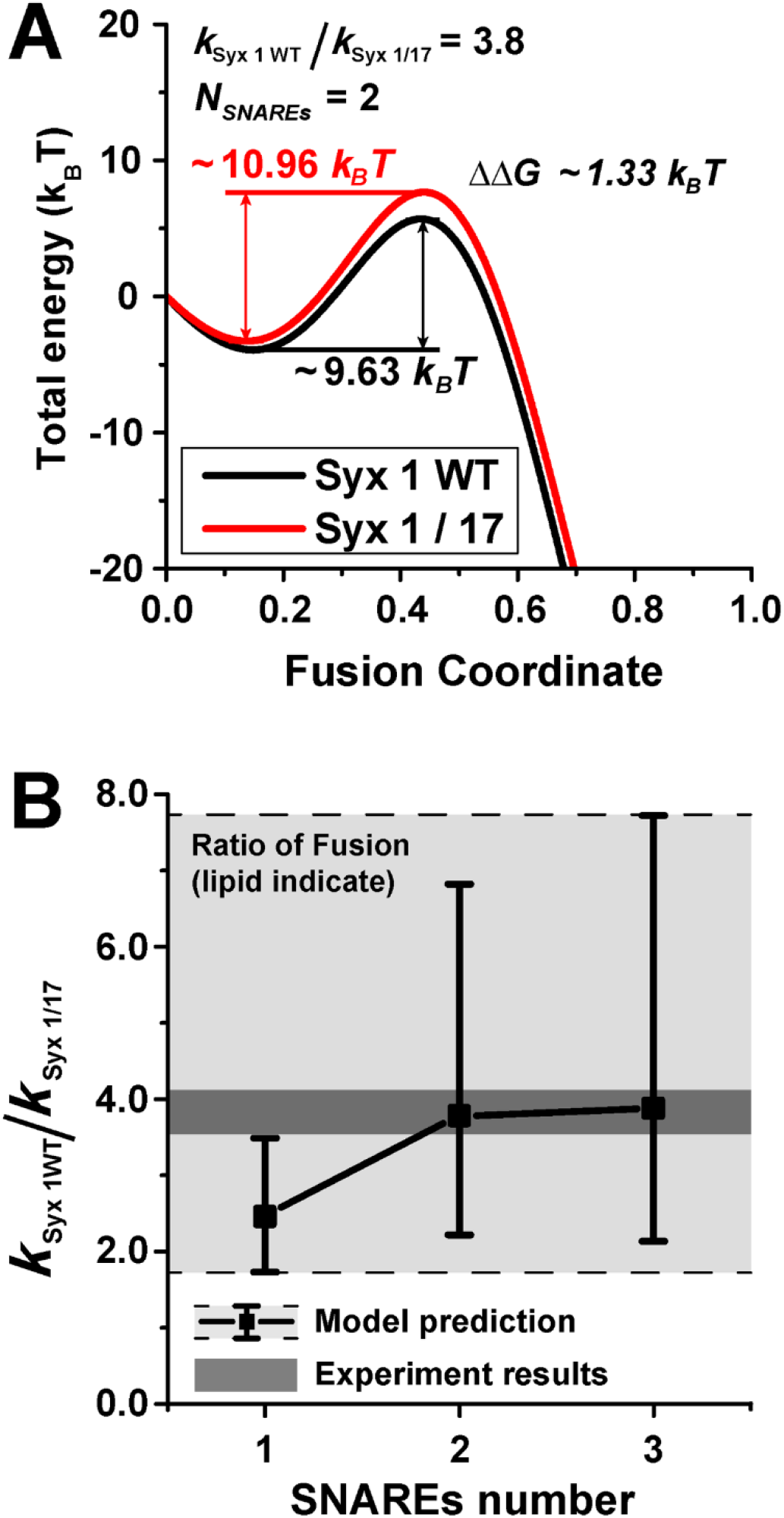
Theoretic analysis of fusion behaviors of systems with Syx 1 WT and Syx 1/17. (A) The energy profiles of Syx 1 WT and Syx 1/17 as function of fusion coordinate. Two SNAREs were assumed to take part in the fusion process. (B) The ratio of fusion rate between Syx 1 WT and Syx 1/17 with various SNAREs number predicted by the theoretical model that was compared with the experiment results. The solid line with square symbol illustrates that tilt energy *E*_t0_ = 2.25*k*_*B*_*T* (20% Chol) and SNAREs number varies from 1 to 3. The light grey shadow region illustrates that tilt energy *E*_t0_ varies from 1.4 to 3.1 *k*_*B*_*T* (20% Chol) and SNAREs number varies from 1 to 3.

## DISCUSSION AND CONCLUSIONS

The interplay between the protein and membrane is essential for the membrane fusion [53]. As the direct link between SNAREs and membrane, the TMD stays the center of biophysical and biochemical researches of SNARE-mediated membrane fusion [54] [55]. Here in this study, we studied how the structure of TMD influenced the fusion rate with the development of a unique technique of adding an extra TMD in the protein. We showed that the dTMDs led to a slower membrane fusion rare compared with the sTMD. The mechanism is that the dTMDs induced larger protein-lipid mismatch. Both of our experimental and theoretical results showed that the protein-lipid mismatch could significantly increase the energy barrier for the membrane fusion and thus reduces the fusion rate.

Furthermore, we analyzed how the number of SNAREs influences the fusion rate. The SNARE number involved in a fusion process is a hotspot in the fusion related researches [13, 50, 56-59]. The controversy regarding the number still exists in the field, and the proposed number range from one to some dozens [57]. Our results showed that 2-3 SNAREs are more likely involved in a general fusion process, consistent with previous studies [50, 57-59] that reported that one to three SNAREs are sufficient for completing the membrane fusion [50, 59]. This study provides a quantitative understanding on the effect of structure of SNAREs on the regulation of the membrane fusion.

## Author Contributions

D.L., B.J., and J.D. contributed to concept and design of this study. B.B. performed theoretical calculation. Z.T. performed and analyzed data for fusion and docking experiments. B.B., D.L., K.Z., W.C., B.J., and J.D. wrote manuscript.

## Acknowledgments

The authors would like to thank Mark Padolina in Axel Brunger’s Lab at Stanford University for protein preparation. K.Z. was supported by the School of Molecular Cell Biology at the University of Illinois at Urbana-Champaign. J.D. was supported by the NIH (R35GM128837 to J.D.).

## Notes

### Competing Interest Statement

The authors have declared no competing interest.

### Summary of Updates

One new author; All figures revised; main text revised

