## Supporting Figures for "Double-transmembrane domain of SNAREs decelerates the fusion by increasing the protein-lipid mismatch"

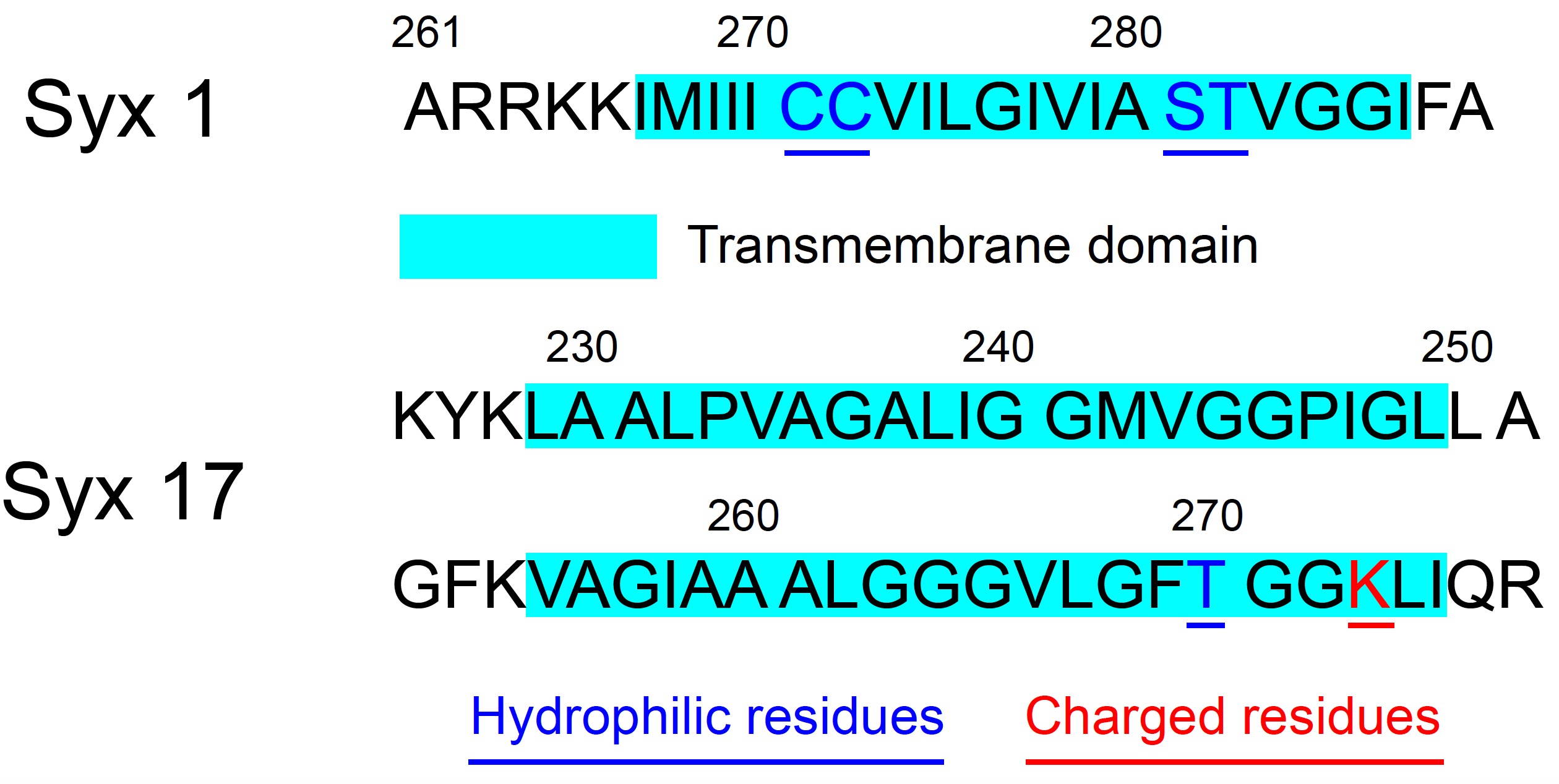

**S1 Fig.** The residues sequences of Syx 1 and Syx 17 TMDs.

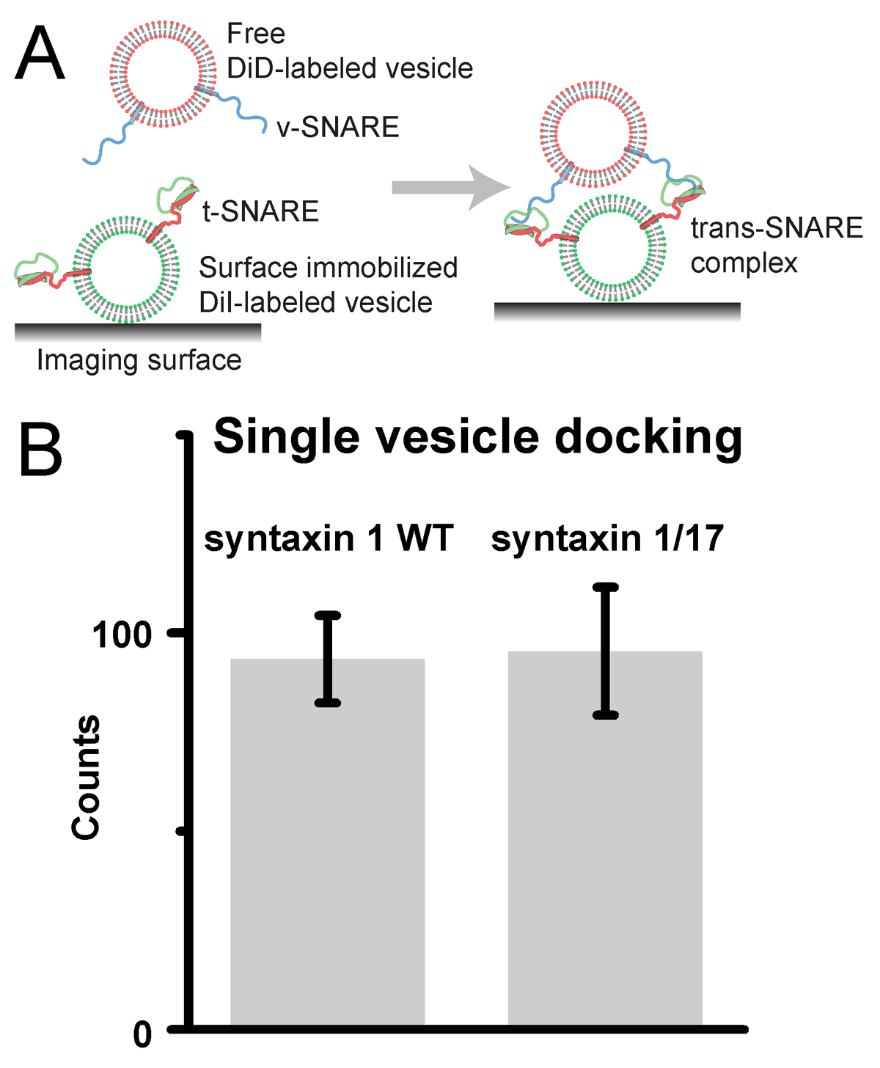

**S2 Fig.** The double TMD of syntaxin 1/17 has little influences vesicle docking in vitro. (A) The illustration of v-vesicles and t-vesicles docking. (B) Quantification of single-vesicle docking assay. Error bars represent SD.

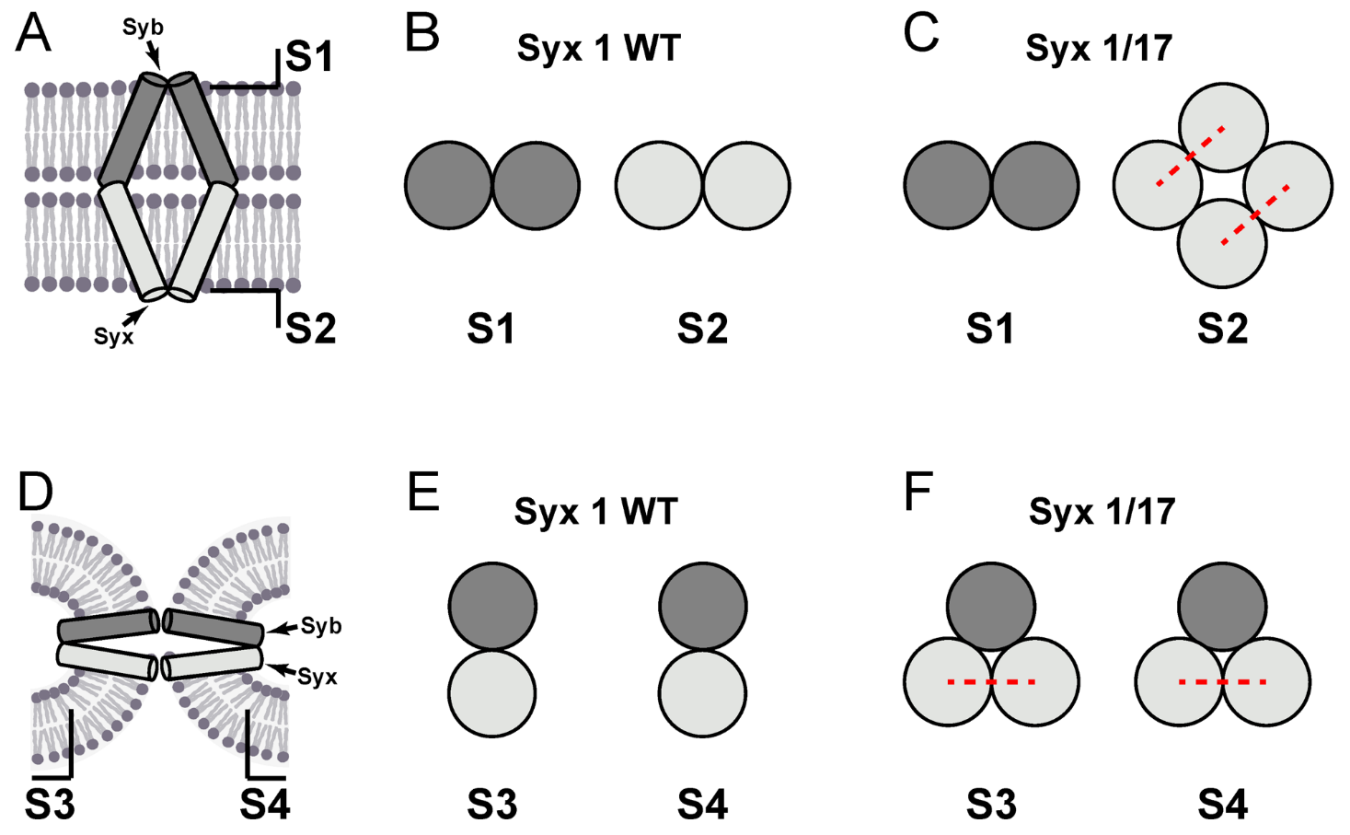

**S3 Fig.** Illustration of the differences for the lipid-protein mismatch with Syx 1 WT and Syx 1/17. (A) Sideview of the structure of TMDs and membranes before fusion. The symbols S1 and S2 indicate the position of sections in (B) and (C). (B) When two SNAREs were involved with Syx 1 WT, there are two Syb TMDs in section S1 and two Syx TMDs in section S2,
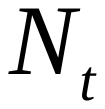
 comes to four. (C) When two SNAREs were involved with Syx 1/17, there are two Syb TMDs in section S1 and four Syx TMDs in section S2,
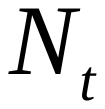
 comes to six. (D) Sideview of the structure of TMDs and membranes after fusion. The symbols S3 and S4 indicate the position of sections in (E) and (F). (E) When two SNAREs were involved with Syx 1 WT, the TMDs of Syb and Syx 1 WT came to close contact with each other in section S3 and S4. (F) When two SNAREs were involved with Syx 1/17, the single TMD of Syb came to contact with the double TMDs of Syx 1/17 in section S3 and S4.
